# Role of EnvC and the phosphoenolpyruvate:sugar phosphotransferase system in resistance to MreB disruption

**DOI:** 10.1101/2022.07.18.500564

**Authors:** Ryan Sloan, Jacob Surber, Emma Roy, Ethan Hartig, Randy M. Morgenstein

**Affiliations:** Department of Microbiology and Molecular Genetics, Oklahoma State University, Stillwater Ok

**Keywords:** MreB, A22, elongasome, divisome, phosphoenolpyruvate:sugar phosphotransferase system, cell shape

## Abstract

Cell wall synthesis in bacteria is determine by two protein complexes: the elongasome and divisome. The elongasome is coordinated by the actin homolog MreB while the divisome is organized by the tubulin homolog FtsZ. While these two systems must coordinate with each other to ensure that elongation and division are coregulated, this cross talk has been understudied. Using the MreB depolymerizing agent, A22, we found that multiple gene deletions result in cells exhibiting increased sensitivity to MreB depolymerization. One of those genes encodes for EnvC, a part of the divisome that is responsible for splitting daughter cells after the completion of cytokinesis through the activation of specific amidases. Here we show this increased sensitivity to A22 works through two known amidase targets of EnvC: AmiA and AmiB. In addition, suppressor analysis revealed that mutations in the phosphoenolpyruvate:sugar phosphotransferase system (PTS) can suppress the effects of A22 in both wild-type and *envC* deletion cells. Together this work helps to link elongation, division, and metabolism.

## Introduction

Bacteria come in a variety of shapes and sizes that are determined by the peptidoglycan cell wall. One of the more common shapes is a bacillus, or rod shape. Rod shape is regulated by the actin homolog, MreB. In addition to MreB many proteins are needed to form and maintain a rod shape. Together these proteins are termed the elongasome and are made of penicillin binding proteins (PBP), SEDS proteins (shape, elongation, division, and sporulation) and different accessory factors, such as MreC, MreD, and RodZ (1-8). MreB is thought to act as a scaffold to organize and direct the localization of the cell wall synthesis enzymes to maintain rod shape (9-12).

MreB is highly conserved among bacteria and can be found in both Gram-positive and Gram-negative species. MreB forms short polymers on the inner surface of the cytoplasmic membrane (8, 13, 14). Loss of *mreB* leads to spherical cells that eventually lyse, unless grown very slowly or with suppressor mutations, such as the overexpression of the cell division genes, *ftsZAQ* (15). To avoid accumulating suppressor mutations, the MreB depolymerizing agent A22 is often used to study the role of MreB in cell shape and physiology (16, 17).

We have previously explored conditions in which *Escherichia coli* cells are better able to withstand the loss of MreB function. Using A22 as a method for disrupting MreB we have shown that MreB and the elongasome in general are dispensable for growth but not cell shape once cells reach a threshold density (18). In addition, we have shown that mutations in metabolic genes that lead to an increase in cell wall precursors allow cells to better tolerate A22 treatment (19). To further understand the role of MreB in cell physiology we sought to find mutants that lead to an increase in sensitivity to A22 treatment.

Here, we used the Keio collection to find mutants that grow poorly in low levels of A22 and found that deletion of five genes result in reduced growth when MreB is disrupted (20). One of these genes, *envC*, is an activator of the cell wall amidases, AmiA and AmiB, which are needed for cell separation after division (21, 22). EnvC is part of the divisome, a separate cell wall synthesis complex used during cell division that is distinct from the MreB elongasome (23). We show that the increased A22 sensitivity is due to the lack of AmiA and AmiB activity and can be suppressed by mutations in MreB or deletion of enzyme 1 of the phosphoenolpyruvate:sugar phosphotransferase system (PTS). This work further connects central metabolism and cell size.

## Results

### Deletions of diverse genes lead to A22 sensitivity

MreB is an essential gene for determining rod shape in many bacteria. It is conditionally essential for viability, as mutations that upregulate the cell division genes *ftsZAQ* or very slow growth can suppress the lethality of *mreB* deletions without restoring rod shape (15). It is unknown if there are genetic conditions that make cells more susceptible to disruption of MreB. It is also unclear if MreB plays other roles in the cell, independent from regulating rod shape. We have previously reported that two gene deletions, *envC* and *tusE*, result in cells with increased sensitivity to A22, although the mechanism for this is unknown (18). Interestingly, deletion of either of these genes leads to slower growth, a condition normally thought to promote growth without MreB (Fig S1) (18).

In order to determine if MreB has other cellular roles we performed a screen of the Keio collection, an ordered library of *E. coli* mutants, to find strains that are more sensitive to growth in low levels of A22, an MreB depolymerizing drug (16, 17, 20). The Keio collection was grown in either LB medium or LB supplemented with a non-lethal amount of A22 (1 µg/ml). At this concentration of A22, wild-type (WT) cells grow close to the same rate as in LB alone; therefore, we looked for deletion strains that have reduced growth in A22 compared to LB alone to determine which, if any, gene deletions lead to increased sensitivity to MreB disruption. After three trials, any deletion that had reduced growth in A22 compared to LB in more than one trial was transduced from the Keio collection into MG1655, the WT *E. coli* strain used in our lab.

Five gene deletions were found to have an increased sensitivity to A22 (Table S1, Fig.1A). To further characterize this sensitivity, these strains were grown in low (1 µg/ml) or high (10 µg/ml) levels of A22 for six hours. As the different mutant strains may have reduced growth rate in LB alone, the O.D._600_ of cells grown in A22 was compared to growth in LB medium alone to produce a growth ratio. In this way we can measure the specific effect of A22 on growth for each strain. WT MG1655 cells grow to nearly the same density in six hours in LB or low A22, producing a growth ratio ∼0.9, but have about a 50% reduction in growth when grown in high levels of A22 (Fig. 1A). All five strains from our screen display lower growth in high levels of A22 with three stains also having reduced growth in low levels of A22. In support of our findings (*ΔsecB*), defects in the Sec system have been shown to cause increased sensitivity to A22 (24). The *envC* mutant displays the most dramatic phenotype, with almost no growth in high levels of A22.

**Figure 1.**
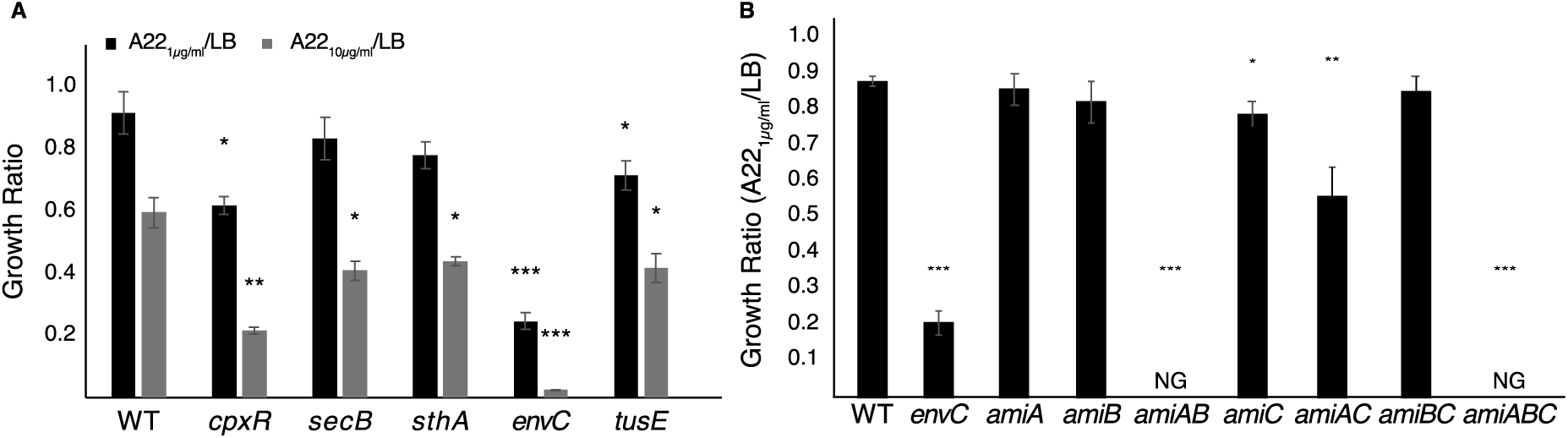
A22 sensitivity is increased in multiple gene deletions. Cells were grown for six hours in LB or LB + A22. The ratio of O.D._600_ between the two conditions was determined as a growth ratio. A) The growth ratio was determined for WT (MG1655) and five gene deletions found through a screen of the Keio collection. Mutations were transduced into MG1655. B) Growth ratios were determined for different strains related to cell separation after division. A-B) Error bars are standard deviation from three independent trials. * p <0.05, ** p <0.01, *** p <0.001

EnvC is an activator of cell wall hydrolases that are needed to separate daughter cells after division (21, 25). Due to the role of EnvC in cell division and therefore possible connection to MreB and the severe growth defect seen in both low and high levels of A22, we focused on determining why deletion of this gene leads to increased sensitivity to MreB disruption. To confirm that deletion of *envC* is responsible for the observed A22 sensitivity, we complemented *envC* on an arabinose inducible plasmid. Induction of plasmid-born *envC* fixes the growth defect in A22 (Fig. S1).

### A22 sensitivity in the *envC* mutant is caused by the loss of AmiAB activity

EnvC works by activating two cell wall hydrolases, AmiA and AmiB (21). To determine if the A22 sensitivity of the *envC* mutant is due to the lack of activation of AmiA and/or AmiB we made deletions of both genes individually and together. Deletion of either *amiA* or *amiB* alone has no effect on sensitivity to low levels of A22 (Fig. 1B). However, the double deletion strain is even more sensitive to A22 than the *envC* deletion and did not grow at low levels of A22 (Fig. 1B). This suggests that the sensitivity of the *envC* mutant to A22 is due to the lack of AmiAB activation and not a secondary role of EnvC. AmiA and AmiB are not the only cell wall hydrolases in *E. coli*. AmiC is a third hydrolase that is activated by the protein NlpD (21, 22). Because loss of *amiA* and *amiB* causes cells to become sensitive to A22, we tested whether loss of AmiC has any effect. Cells lacking AmiC show a mild increase in sensitivity to A22 (Fig. 1B). We further analyzed double mutants of *amiAC* and *amiBC* as well as the triple *amiABC* mutant for A22 sensitivity. The *amiAC* mutant is even more sensitive than the *amiC* mutant alone, but not nearly as sensitive as either the *envC* deletion or *amiAB* mutant. As expected, the triple mutant shows an extreme sensitivity phenotype similar to the double *amiAB* mutant and did not grow in A22 (Fig. 1B). These results suggest that activation of the hydrolases AmiA and AmiB is most important when MreB is disrupted.

As might be expected, the *envC* and *amiAB* deletions result in similar phenotypes of elongated and chained cells (Fig. 2A) (21). As both strains are sensitive to A22 we examined all the strains under the microscope when grown with and without A22 to determine what effect A22 has on cell shape. All cells were grown in LB for two hours before A22 (10 µg/ml) was added and cells were allowed to grow for another two hours. An LB only control was allowed to grow for four hours in total. As expected, WT cells become spherical upon A22 treatment. A similar change in shape was observed for all strains (Fig. 2A). The *amiAB* double deletion exhibits a similar phenotype to the *envC* mutant both with and without A22. Interestingly, the *amiAC* deletion strain exhibits a more pronounced chaining phenotype than the *amiAB* mutant when grown in LB (Fig. 2A) (26). This suggests that the increased sensitivity of the *envC* and *amiAB* mutants to A22 is not due to those cells being chained or elongated, as the *amiAC* strain is more resistant to A22 than either of those strains and forms longer chains (Fig. 1B and 2A).

**Figure 2.**
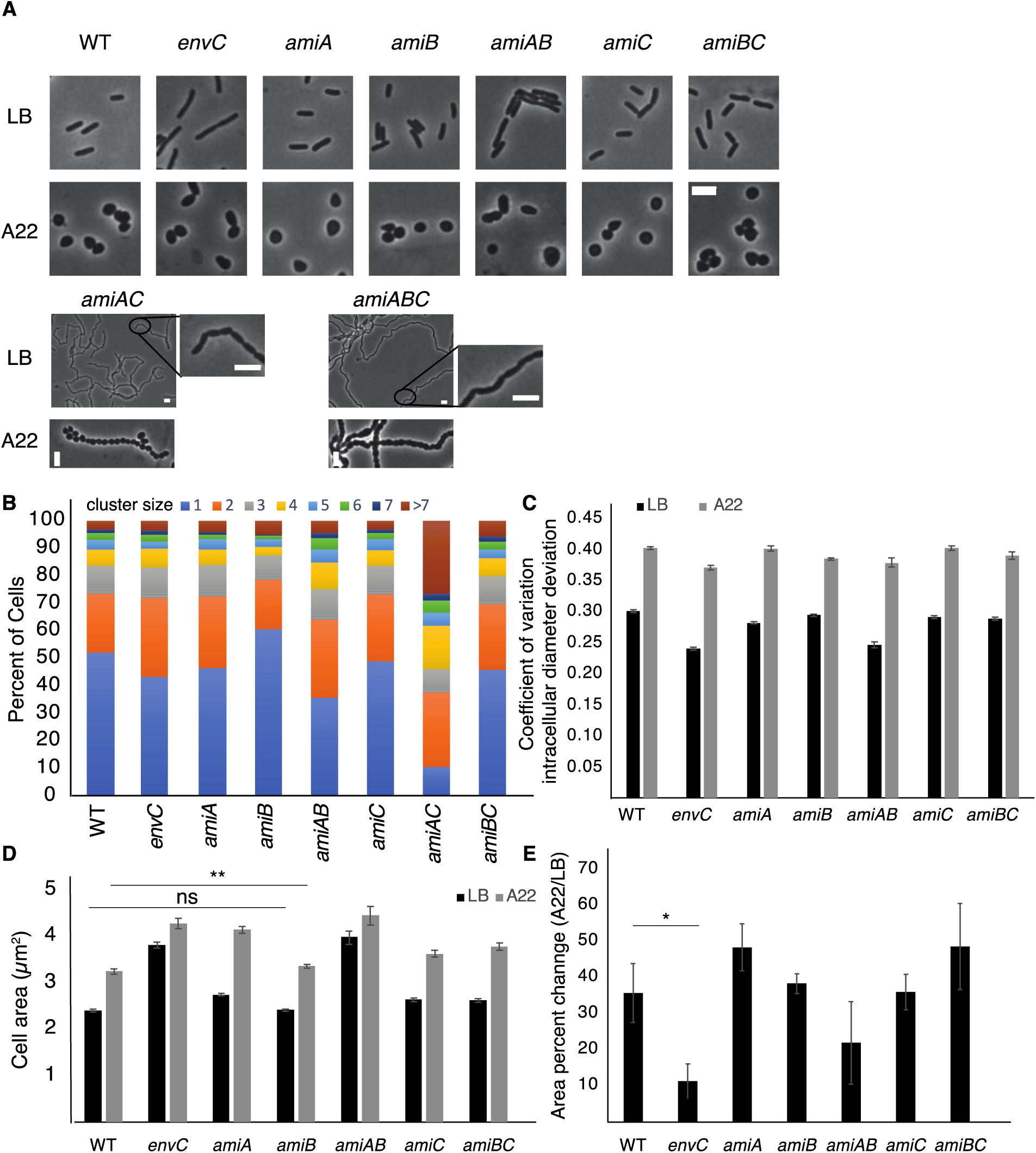
Cell shape characteristics of mutants involved in cell separation with and without A22. A) Cells were grown in LB for four hours or LB for two hours and then A22 (10 µg/ml) for two hours before being imaged. Scale bar = 4µm. B) Cell cluster sizes were counted from images of cells grown in A22 as in A. Shown is pooled data from three independent trials. C-E) Data is pooled from three independent experiments. Cells were grown for 2 hours in LB before the addition of A22 for two hours. C) Coefficient of variation diameter deviation is a metric to measure how ‘rodlike’ cells are. The higher the number the less rod and more spherical a cell is. D) Cell area measurements of cells grown in A. Only individual cells were counted, and chains of cells were ignored in the analysis. All statistical comparisons were made between WT LB and mutants LB or WT A22 and mutants A22 and have a p value < 0.001 unless noted. ** p < 0.01. See table S3 for number of cells. C-D) error bars are 95% CI. E) Percent change of cell area between cells grown in LB or with A22. Error bars are standard deviation from three independent experiments. All comparison are to WT and not significant unless noted. * p <0.5.

### Loss of amidase activity increases sensitivity to envelope targeting antibiotics

It is possible that the above phenotypes are due to a general mechanism increasing cell permeability rather than a specific MreB mechanism. It has been shown that deletion of *envC* causes an increase in permeability as seen by sensitivity to vancomycin and bacitracin, cell wall targeting antibiotics that normally do not pass through the outer membrane (27). This permeability is thought to work through the amidases as mutants in different amidases also lead to increases in cell permeability (27-29).

To test if the *envC* mutant is more sensitive to all antibiotics, we tested for growth in a low and high amount of ampicillin, tetracyline, and chloramphenicol. The low amount is below the inhibitory concentration for WT cells and the high amount shows a significant decrease in grow of WT cells. The *envC* mutant only displays an increased sensitivity to ampicillin, the cell well targeting drug, but not the ribosomal targeting antibiotics (Fig. S2A). These data suggest that loss of amidase activity leads to increased sensitivity to envelope targeting antibiotics in *E. coli*, but not a generalized sensitivity to all antibiotics; therefore, we do not believe cell permeability is the cause of the increased sensitivity.

### Disruption of MreB reduces chaining caused by loss of amidase activity

It is well established that *envC* or amidase mutants form chains of cells when grown in LB (22, 25, 26, 30). This chaining phenotype is not recapitulated in A22. After A22 treatment, WT, *envC*, and *amiAB* strains all have ∼70% of their cells alone or in pairs, with a small percentage of cells found in clusters of larger groups. However, the *amiAC* mutant has ∼30% of its cells found in large (>7 cells) clusters (Fig. 2B). These data suggest some crosstalk between MreB and the cell separation machinery that requires AmiA. When MreB is polymerized (LB), the lack of AmiA and AmiB leads to chained cells, but when MreB is depolymerized (A22 treatment) there is no additional chaining. However, in either a polymerized or depolymerized state of MreB cells that lack AmiA and AmiC form chains. This suggests that the loss of AmiA in parallel with the loss of another amidase produces different phenotypes depending on the polymerization state of MreB, but if AmiA is functional chaining is reduced independent of the state of MreB, explaining why there is no chaining in an *amiBC* mutant.

We further characterized the effects of A22 on each of these mutants by measuring the change in rod shape and size of the cells. Using the metric, coefficient of variation of intracellular diameter deviation (IDD), we can measure the “rodness” of cells (4, 12, 18).

A centerline is drawn through the long axis of the cell and the diameter is measured across the cell body. The change in the standard deviation of these diameters is used to determine “rodness”, as a sphere will have a greater standard deviation than a rod. All strains tested become spherical upon A22 treatment (increased IDD) (Fig. 2C). As cell length will affect the IDD value (longer cells reduce the influence of the poles) and *envC* cells are longer (Fig. S3) we did not attempt to statistically compare IDD values across strains. Not only do all cells treated with A22 become spherical, but there is a concurrent increase in cell area (Fig. 2D). Interestingly, the increase in area upon A22 treatment is smaller in Δ*envC* cells than WT cells (Fig. 2E).

### A22 sensitivity caused by loss of *cpxR* is not due to changes in amidase expression

The CpxR/CpxA two-component system has been shown to positively regulate the expression of both *amiA* and *amiC* (31). As a mutant of *cpxR* was found to be more sensitive to A22 (Fig. 1A) and an *amiAC* double mutant is more sensitive to A22 (Fig. 1B) we asked if the sensitivity of the Δ*cpxR* deletion strain was due to loss of *amiA* and/or *amiC* expression. We cloned *amiA, amiB*, and *amiC* into the vector pTrc99A under the control of a leaky lac promoter and attempted to suppress the A22 sensitivity of the Δ*cpxR* strain through ectopic expression of any of the amidases. While there does appear to be some effect of the empty vector on A22 sensitivity, expression of any amidase was unable to restore growth in A22 to WT levels in the *cpxR* mutant (Fig. S4). This suggests a distinct mechanism of A22 sensitivity in the *ΔcpxR* strain.

### Suppressor mutations in *the envC* deletion strain can grow in A22

We have shown that an *envC* deletion is more sensitive to A22 treatment in an AmiAB dependent manner; however, it is still unclear why the loss of AmiAB function results in increased A22 sensitivity. The *envC* mutant does not grow in high levels of A22, whereas WT cells are able to grow (Fig. 1). We performed a random suppressor screen, by plating the *envC* mutant on A22 plates to find mutants that restore growth. Colonies that grew on A22 plates were streaked onto fresh A22 plates before being grown in liquid cultures with A22, at 37°C. These experiments were performed independently three times. As a control of the acquisition of non-allele specific mutations that make the cells resistant to A22, we moved a known A22 resistant MreB point mutation (MreB_S14A_) into the *envC* deletion (12). This resistant *envC* strain is able to grow in A22 and has an elongated/chained phenotype when grown in LB, although the poles appear to taper. Any colony that grew throughout this selection process was tested for its A22 growth ratio in low and high levels of A22.

We classified the suppressor mutations into three categories based on A22 resistance. Group 1 consists of four mutants (#s 2, 3, 5, 27) that display a high level of resistance, most similar to the WT or Δ*envCmreB*_*S14A*_ strains. Group 2 mutants (#s 7, 15) do not show much change in low levels of A22 but are able to grow in A22 10 µg/ml, unlike the parental *envC* deletion. Group 3 (#s 8, 10, 38) consists of mutants that grow very well in low A22 with only show a small increase in growth at higher A22 levels (Fig. 3).

**Figure 3.**
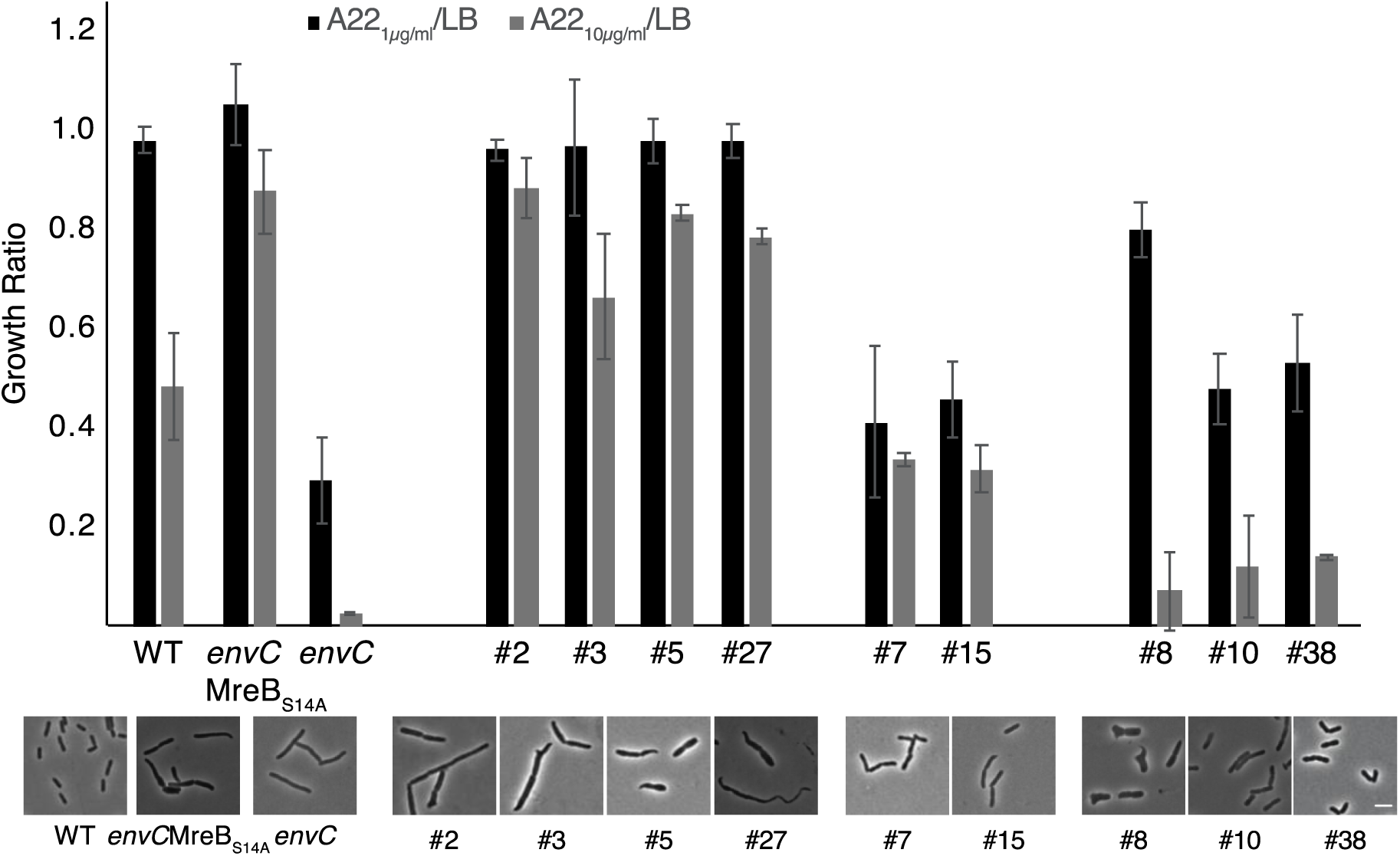
Suppressor screen for mutants of Δ*envC* cells that can grow in A22. Suppressors were selected with multiple rounds of growth on A22 10 µg/ml. Top-Cells are separated into four groups. The leftmost is a group of control strains, including WT, Δ*envC*, and Δ*envC* + an MreB point mutation resistant to A22 (MreB_S14A_). Group 1 contains four strains with high levels of A22 resistance. Group 2 has two strains with increased growth in high levels of A22 and group 3 has three strains with high growth in low A22 and increased growth in high levels. Error bars are standard deviation from three independent experiments. Bottom-representative images of cells grown in LB medium. Scale bar = 4 µm.

To help determine the function of these suppressor mutations we imaged all strains in LB to compare their cell shapes with WT, *ΔenvC*, and *ΔenvCmreB*_*S14A*_ stains, and sequenced the MreB. Five of the nine suppressors have MreB point mutations (Table S2). Interestingly, all group 1 mutants have MreB point mutations, although mutants #5 and #27 have unique shapes that deviate from the shape of the *envC* mutant, suggesting that these mutations have a function distinct from generic A22 resistance. The other MreB point mutant is in group 3 (#8) and results in loss of the elongation and chaining seen when *envC* is removed. This mutant and mutant #27 have a missense mutation in residues close to each other yet have different A22 resistance profiles (Table S2).

We performed whole-genome sequencing on all mutants without an MreB mutation (#s 7, 15, 10, 38) and the three mutants with MreB point mutations with shapes that deviate from the MreB_S14A_ control (#s 5, 27) or are in a different resistance group (#8) (Table S2). Due to their similar shape and resistance profile to *ΔenvCmreB*_*S14A*_, we did not perform whole-genome sequencing on #s 2 and 3.

Both strains in group 2 have an insertion in *ptsI*, encoding for enzyme 1 of the PTS (32). The three mutants in group 3 have no mutation in common. Mutant #8 has an MreB point mutation, while the sequencing of mutant #38 did not reveal any changes. Mutant #10 has a ∼13kb deletion of many genes involved in chemotaxis and flagella rotation. Due to known connections between metabolism and cell shape we will focus on the role of *ptsI* on A22 resistance (19, 33-35). It is of note that mutants #3 and #5 have mutations in adjacent residues of MreB which are predicted to be in the A22 binding pocket, yet produce different cell shapes (Table S2) (36). As this screen was not saturating it is possible that mutations in other genes can suppress Δ*envC* growth on A22.

### Loss of functional PtsI suppresses A22 sensitivity to Δ*envC*

*ptsI* encodes for enzyme 1 in the PTS, which is used to transport and phosphorylate sugars, such as glucose, as they enter the cell (37). As all the described experiments were performed in LB without the addition of any sugar, it is surprising to find suppressor mutations in *ptsI*. The suppressors have the same two base pair insertion in the gene; therefore, we tested whether a deletion of *ptsI* also increases A22 resistance. Deletion of *ptsI* in a WT background leads to increased resistance to high levels of A22 and can recapitulate the suppressor phenotype in an Δ*envC* background (Fig. 4A), supporting the hypothesis that the original suppressor phenotype came about from the loss of *ptsI*.

**Figure 4.**
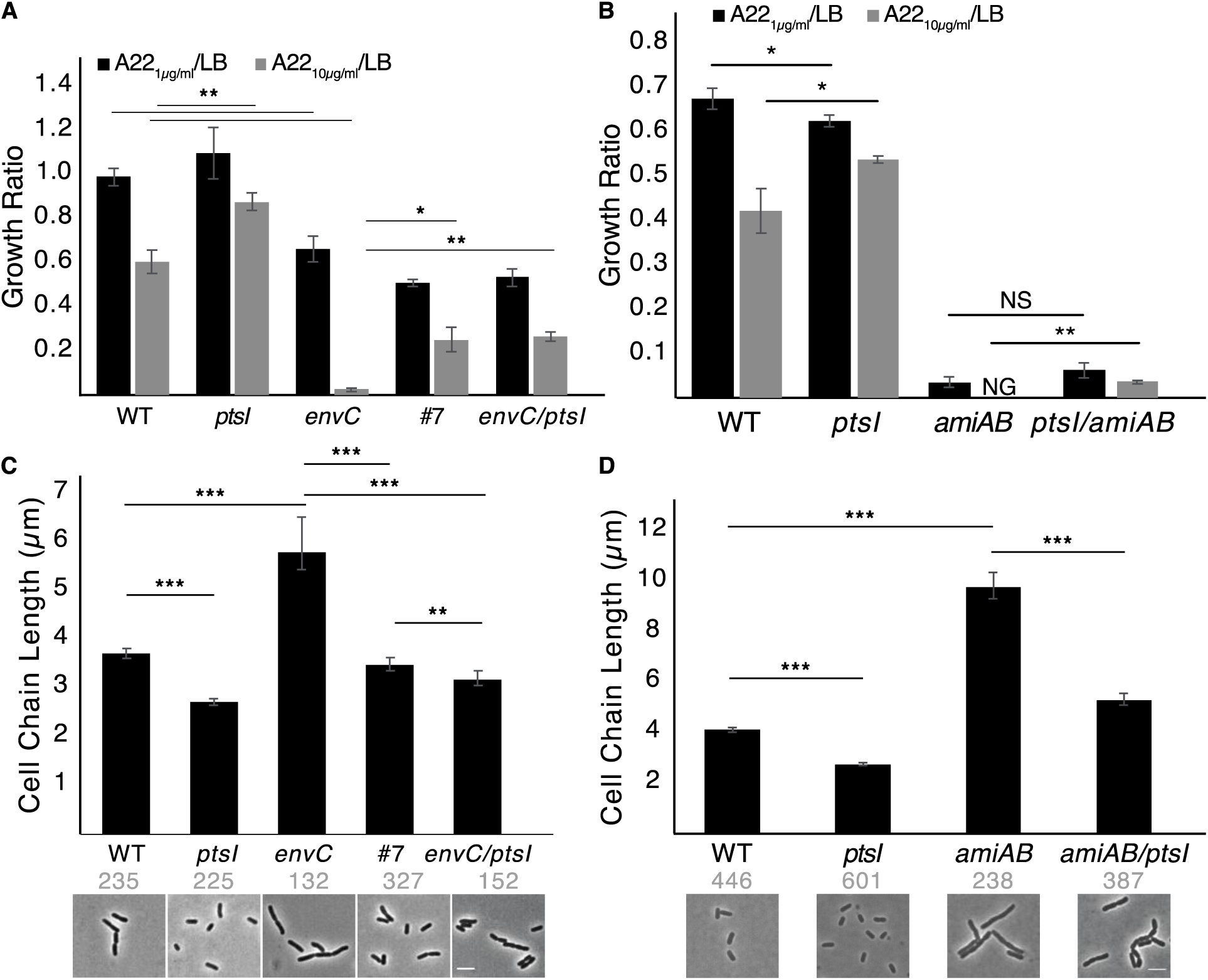
Deletion of *ptsI* suppresses both A22 and cell elongation phenotypes of Δ*envC* cells. A-B) Average growth ratio from three experiments of cells grown in LB or LB + A22. Error bars are standard deviation. * p <0.05, ** p <0.01. C-D) Cell length of WT and indicated mutant cells. Cells in chains were included to capture the effect of chaining in different backgrounds. Error bars are 95% CI and scale bar = 4 µm. ** p < 0.01, *** p < 0.001.

To further confirm the role of PtsI in A22 resistance we complemented the *ptsI* and *envCptsI* deletion strains with either a WT allele of PtsI or an allele that in unable to be phosphorylated (PtsI_H189A_) (38, 39). If PtsI phosphorylation is linked to its role in A22 sensitivity than the H189A mutant should mimic the deletion strain and provide increased resistance to A22. We found that complementation with the WT allele makes both the *ptsI* and *envCptsI* deletion strains more sensitive to A22 than when complemented with the nonphosphorylatable allele (Fig. S5), supporting our hypothesis that the loss of PtsI activity increases the resistance of cells to disruption of MreB.

One of the common mechanisms for suppression of the lethality of *mreB* disruption is the upregulation of the *ftsZAQ* operon. It is possible that deletion of *ptsI* leads to increased levels of the *ftsZ* operon providing resistance to A22. We transduced a functional fluorescent copy of FtsZ into the native site of the chromosome as the sole copy of FtsZ in the cell (19). A *ptsI* deletion strain expressing this construct is still smaller than WT cells (Fig. S6). These strains allowed us to use fluorescence intensity to determine if there are changes in FtsZ levels caused by deleting *ptsI*. We found similar albeit slightly lower levels of FtsZ-GFP in the Δ*ptsI* strain when accounting for cell size (3.88 × 10^5^ ± 2.77 × 10^5^ versus 3.03 × 10^5^ ± 2.25 × 10^5^ AU/pixel) suggesting that the suppressive effect of *ptsI* deletion is not due to increases in *ftsZ* levels. We also noticed that cells deleted for *ptsI* are more likely to lack an FtsZ structure. Although this was rare in both WT (4.5%) and *ΔptsI* (8.5%) cells the lack of FtsZ structures would not account for smaller cells (Fig. S6 white arrows).

In addition to A22, we found that Δ*envC* cells are sensitive to ampicillin. To determine if the deletion of *ptsI* is specific to A22 treatment or more general cell wall stresses, we tested the sensitivity of the *envCptsI* double mutant in ampicillin and found that the loss of *ptsI* is able to suppress the sensitivity to ampicillin at high concentrations, similar to that seen with A22 (Fig. S2B).

### Loss of *ptsI* suppresses A22 sensitivity to Δ*envC* but not Δ*amiAB* cells

We have shown that the sensitivity to A22 seen in *envC* cells works through *amiAB* (Fig. 1B); therefore, we tested if deletion of *ptsI* would also be able to suppress the A22 sensitivity in an *amiAB* background. Interestingly, while deletion of *ptsI* in an *envC* background leads to an 11X fold increase in growth ratio in A22 10 µg/ml, in an *amiAB* deletion background there is only a 0.02X fold change and growth is most likely below the accuracy level of the spectrophotometer (Fig. 4AB). This suggests that the suppressive effect of losing PtsI works through AmiA or AmiB. This datum also suggests that the loss of PtsI does not change membrane permeability, as we would expect all strains that have increased A22 sensitivity to become more resistant if permeability of A22 was affected.

### Loss of *ptsI* suppresses the cell elongation phenotype of an *envC* mutant

One of the initial observations of an *envC* mutant was that it forms elongated and chained cells (Fig. 3A, Fig. 4C, Fig. S3) (25). Because deletion of *ptsI* suppresses the A22 sensitivity phenotype through AmiA/AmiB, we asked whether cell size is also affected. Cells were grown in LB without additional sugar, to mid log phase and imaged. To account for the chaining effect of the *envC* mutant, we measured both individual and chained cells. As expected, Δ*envC* cells are longer than WT; interestingly, the lack of *ptsI* reduces the cell length of WT cells and Δ*envC* cells (Fig. 4C). Since deletion of *ptsI* is unable to provide *amiAB* cells A22 resistance, we examined if deletion of *ptsI* would reduce the cell size of the *amiAB* mutant. Deletion of *ptsI* is able to reduce cell length in the absence of *amiAB* (Fig. 4D). These results show that loss of *ptsI* can suppress both the A22 sensitivity and cell length phenotype resulting from the loss of *envC*, but only the cell length phenotype of an *amiAB* mutant, suggesting that the role of PtsI in A22 resistance can be separated from its role in cell size regulation and supporting a role for AmiAB in A22 sensitivity.

## Discussion

MreB is a highly conserved protein essential for the regulation of rod shape and viability, making it a potentially good target for antibiotic development (1, 15). While we have previously reported on conditions that help cells grow without MreB, there is little known about how cells can become hypersensitive to the disruption of MreB (18, 19). Here, we performed a screen of the Keio collection to identify gene deletions that lead to the inhibition of growth in sublethal levels of A22, an MreB depolymerizing agent. Five strains were found to have increased sensitivity to A22, one of which, *envC*, is involved in cell division (Fig. 1A).

There are two distinct cell wall synthesis complexes in rod-shaped bacteria, the MreB-regulated elongasome controlling cell elongation and growth, and the FtsZ-regulated divisome controlling cell division. If and how these two systems talk with each other is underexplored. While it is well established that MreB localizes to the Z-ring during division in *Caulobacter crescentus*, it is more controversial in *E. coli* (40, 41). While our lab and others have not been able to see an enrichment of MreB at the division site when using a functional fluorescent sandwich fusion of MreB as the only copy in the cell, others have reported that N-terminal fusions to MreB and the MreB interaction partner, RodZ, localize to the midcell (42-44). This study shows that there is some crosstalk between these two systems as loss of *envC* (divisome) makes cells more sensitive to disruption of MreB (elongasome) with A22.

The “adder-model” of cell size regulation says that rather than reach a critical size before dividing, cells add a constant volume before dividing (45, 46). It would reason that there is coordination between lateral wall synthesis controlled by MreB and cell division. Why this system would work at the level of cell separation after division is unclear, but it may suggest the role of proteins upstream or downstream of EnvC, such as FtsN (25, 47).

### Loss of *amiA* and *amiB* phenocopy *envC’s* A22 sensitivity

EnvC activates two cell wall hydrolases, AmiA and AmiB, needed to separate daughter cells after division has occurred. To determine if the sensitivity of the *envC* deletion is due to a secondary role of EnvC or its role in activating AmiA and AmiB, we deleted both proteins alone and together. The double *amiAB* deletion was even more sensitive to A22 than the *envC* deletion (Fig. 1). This is most likely due to latent activity of either enzyme in the *envC* deletion that is absent when the proteins are deleted. It will be interesting in future work to determine if AmiA and/or AmiB have a secondary role in the cell that when absent leads to cells being more sensitive to changes in MreB polymerization.

The activity of the amidases during cell division produces denuded glycan chains (missing peptide side chains) which signal to activate FtsN (48, 49). When active. FtsN activates PBP1B at the division site (50). PBP1B mutants are sensitive to A22, and PBP1B is necessary to overcome the loss of the elongation machinery (18, 19, 51). One possible reason for the increased sensitivity of an *envC* mutant is a lowering of the presence of denuded glycans leading to less active FtsN and therefore less active PBP1B. Without a highly active PBP1B the cell would not be able to overcome the loss of the elongasome.

### Point mutations in *mreB* suppress effects of *envC*

In order to further understand the connection between MreB and EnvC we performed an A22 resistance suppressor screen. Of the nine suppressor strains found in this screen, five have mutations in MreB. Interestingly, while 4/5 of the *envCmreB* mutants have increased A22 resistance over WT cells, they also have unique cell shapes in LB medium (Fig. 3). Mutants #5 and #27 are both shorter than the *envC* parent strain with tapered tails and do not appear to chain. Furthermore, #3 and #5 have mutations in adjacent residues that are predicted to interact with ATP, yet cause different cell shapes (36). Molecular dynamic simulations suggest that ATP binding of the MreB monomer is necessary for polymerization and that polymerization induces structural changes in MreB leading to hydrolysis; therefore, we suggest that these two mutations may help stabilize ATP-binding leading to increased filamentation and A22 resistance (52). Alternatively, these mutations may decrease the exchange between ADP and ATP, stabilizing filaments and leading to A22 resistance. The nucleotide-bound state of MreB polymers may also affect membrane binding or protein interactions, as seen with eukaryotic actin (53, 54).

Residues 72 (#8) and 79 (#27) are near each other on the MreB protein but are not predicted to have a structural role in filamentation and lead to different A22 resistance profiles (Fig. S7). These residues may represent an MreB-protein interaction site. It will be interesting to study the effects of these MreB mutations in a background with EnvC to see if cell shape effects are due to the loss of *envC*.

### PtsI regulates cell size even without excess sugars

Cells can sense their metabolic environment and adjust their size accordingly. The fact that cells grow larger in rich medium and smaller in nutrient poor medium is termed the “Growth Law” (55). One mechanism for connecting nutrient levels with cells size links the inhibition of FtsZ with UDP-glucose levels (56, 57). Our experiments reveal that deletion of *ptsI* results in shorter cells in WT, Δ*envC*, and Δ*amiAB* backgrounds (Fig.5C). Interestingly, these cells were grown in LB medium without the addition of sugar. As the PTS system is involved in sugar uptake, it is surprising to find such a dramatic change in cell length in these conditions.

PtsI is the first protein in the PTS which transfers a phosphate from phosphoenolpyruvate (PEP) and shuttles it to enzyme 2 of the PTS, where it can be added to sugars upon entry into the cell. Lack of PtsI will affect the PEP:pyruvate ratio of the cell (58, 59). We have previously proposed that increased pyruvate levels in cells can activate gluconeogenesis and increase the levels of cell wall precursors providing cells with increased tolerance to A22 (19). In *Bacillus subtilis*, deletion of pyruvate kinase (*pyk*) effects Z-ring formation (60). We propose that deletion of *ptsI* results in a change in the PEP:pyruvate ratio in the cell, resulting in increased resistance to A22 and changes in cell size. This mechanism appears to connect with AmiAB as the deletion of *ptsI* does not result in A22 resistance in this strain. However, it is also possible that changes in the PTS affect cAMP levels, which has been shown to regulate cell shape (35).

## Experimental Procedures

### Bacterial Growth

Bacteria were grown using standard laboratory conditions. Cultures were grown overnight in LB medium (10 g L^-1^ NaCl, 10 g L^-1^ tryptone, 5 g L^-1^ yeast extract) and subcultured in the morning 1:1000 and grown to exponential phase (O.D._600_ 0.2-0.5) at 37°C in a shaking incubator. Keio collection mutants were moved in MG1655 using P1 transduction. Phage were propagated in a WT MG1655 strain before being grown with PCR verified mutants from the Keio collection. Tranducing phage were then grown on MG1655 and transductants were selected on kanamycin (30 μg ml^-1^) and confirmed by PCR. See Table S1 for a list of strains used in this study.

### Growth Ratios

Overnight cultures of cells were subcultured 1:1000 into media with 1 or 10 µg ml^-1^, or no A22. Cultures were grown shaking at 37°C for 6 hours before measuring the O.D._600_. Growth ratios were determined for each strain comparing the A22 cultures to the no drug culture. Error bars were determined by taking the standard deviation of the growth ratio across multiple experiments.

### Suppressor Screen

A22 suppressors of the Δ*envC* mutant were isolated by growing Δ*envC* cells overnight in LB media lacking A22. Cells were plated on LB + A22 (10 µg ml^-1^) and incubated for 16 hours at 37°C. Colonies were streaked onto fresh A22 plates and incubated for 16 hours at 37°C. Colonies were then grown in liquid A22 media for 16 hours.

Mutants that grew in liquid media were subjected to growth ratio analysis in A22 1 and 10 µg ml^-1^ and imaged from exponential phase in LB medium. MreB was PCR amplified and sequenced before strains were sent to MiGS for whole-genome Illumina sequencing. Both the *ΔenvC* parent and suppressors were sequenced. Indicated mutations are the only differences. Variants were found using breseq on census/mixed base mode (61). The consensus mutation E-value cutoff and polymorphism E-cutoff was set to 10 with a frequency cutoff of 0.8 and a polymorphism frequency cutoff of 0.2.

### Microscopy

For all imaging, cells were grown at 37°C in LB medium. Imaging was done on 1% M63-glucose agarose pads at room temperature. Phase contrast and fluorescent images were collected on a Nikon Ni-E epifluorescent microscope equipped with a 100X/1.45 NA objective (Nikon), Zyla 4.2 plus cooled sCMOS camera (Andor), and NIS Elements software (Nikon).

Cell area and IDD were calculated using the matlab software Morphometrics (62), and custom software as described previously. Only single non-diving cells were used for analysis unless stated otherwise.

### MreB Mutation Modeling

MreB mutations were modeled using Swiss model server (<u>SWISS-MODEL</u> (expasy.org)) using *Caulobacter crescentus* MreB (4czg.1.pdb) (https://swissmodel.expasy.org/templates/4czg.1) as a template. The mutations were mapped using PymoL Version 2.5.2.

## Acknowledgements

Thanks to Dr. Oma Amster-Choderl for PtsI strains. Thank you to Dr. Elizabeth Ohneck for careful reading of the manuscript. This work was funded by NIH <u>1R15GM129636-01A1</u>.

## References

1. Shi H, Bratton BP, Gitai Z, Huang KC. How to Build a Bacterial Cell: MreB as the Foreman of E. coli Construction. Cell. 2018;172(6):1294–305; PMCID: PMC5846203.

2. Meeske AJ, Riley EP, Robins WP, Uehara T, Mekalanos JJ, Kahne D, Walker S, Kruse AC, Bernhardt TG, Rudner DZ. SEDS Proteins are a Widespread Family of Bacterial Cell Wall Polymerases. Nature. 2016;537(7622):634–8; PMCID: PMC5161649.

3. Cho H, Wivagg CN, Kapoor M, Barry Z, Rohs PDA, Suh H, Marto JA, Garner EC, Bernhardt TG. Bacterial Cell Wall Biogenesis is Mediated by SEDS and PBP Polymerase Families Functioning Semi-Autonomously. Nature Microbiology. 2016;1:16172; PMCID: PMC5030067

4. Morgenstein RM, Bratton BP, Ouzounov N, Nguyen JP, Shaevitz JW, Gitai Z. RodZ links MreB to Cell Wall Synthesis to Mediate MreB Rotation and Robust Morphogenesis. Proc Natl Acad Sci USA. 2015:12510–5. PubMed PMID: 26396257.

5. Bendezu FO, Hale CA, Bernhardt TG, de Boer PAJ. RodZ (YfgA) is Required for Proper Assembly of the MreB Actin Cytoskeleton and Cell Shape in E. coli. EMBO. 2009;28(3):193–204; PMCID: PMC2637328.

6. Alyahya SA, Alexander R, Costa T, Henriques AO, Emonet T, Jacobs-Wagner C. RodZ, a Component of the Bacterial Core Morphogenic Apparatus. Proc Natl Acad Sci USA. 2009;106(4):1239–44. doi: 10.1073/pnas.0810794106; PMCID: PMC2633561.

7. Shiomi D, Sakai M, Niki H. Determination of Bacterial Rod Shape by a Novel Cytoskeletal Membrane Protein. The EMBO Journal. 2008;27(23):3081–91. doi: 10.1038/emboj.2008.234. PubMed PMID: PMC2599877; PMCID: PMC2599877.

8. Kruse T, Bork-Jensen J, Gerdes K. The Morphogenetic MreBCD Proteins of Escherichia coli form an Essential Membrane-Bound Complex. Mol Microbiol. 2005;55(1):78–89. PubMed PMID: 15612918.

9. Billings G, Ouzounov N, Ursell T, Desmarais SM, Shaevitz J, Gitai Z, Huang KC. De Novo Morphogenesis in L-Forms Via Geometric Control of Cell Growth. Mol Microbiol. 2014;93(5):883–96. doi: 10.1111/mmi.12703. PubMed PMID: PMC4459576.

10. Ursell TS, Nguyen J, Monds RD, Colavin A, Billings G, Ouzounov N, Gitai Z, Shaevitz JW, Huang KC. Rod-Like Bacterial Shape is Maintained by Feedback Between Cell Curvature and Cytoskeletal Localization. Proc Natl Acad Sci USA. 2014;111(11):E1025–E34. doi: 10.1073/pnas.1317174111. PubMed PMID: PMC3964057.

11. Lee TK, Tropini C, Hsin J, Desmarais SM, Ursell TS, Gong E, Gitai Z, Monds RD, Huang KC. A Dynamically Assembled Cell Wall Synthesis Machinery Buffers Cell Growth. Proc Natl Acad Sci USA. 2014;111(12):4554–9. doi: 10.1073/pnas.1313826111.

12. Bratton BP, Shaevitz JW, Gitai Z, Morgenstein RM. MreB Polymers and Curvature Localization are Enhanced by RodZ and Predict E. coli’s Cylindrical Uniformity. Nature Communications. 2018;9(1):2797. doi: 10.1038/s41467-018-05186-5; PMCID: PMC6052060

13. Salje J, van den Ent F, de Boer P, Lowe J. Direct Membrane Binding by Bacterial Actin MreB. Mol Cell. 2011;43(3):478–87; PMCID: PMC3163269.

14. Dempwolff F, Reimold C, Reth M, Graumann PL. Bacillus subtilis MreB Orthologs Self-Organize into Filamentous Structures underneath the Cell Membrane in a Heterologous Cell System. PLOS ONE. 2011;6(11):e27035. doi: 10.1371/journal.pone.0027035; PMCID: PMC3206058.

15. Bendezú FO, de Boer PAJ. Conditional Lethality, Division Defects, Membrane Involution, and Endocytosis in mre and mrd Shape Mutants of Escherichia coli. J Bacteriol. 2008;190(5):1792–811. doi: 10.1128/jb.01322-07; PMCID: PMC2258658

16. Bean GJ, Flickinger ST, Westler WM, McCully ME, Sept D, Weibel DB, Amann KJ. A22 Disrupts the Bacterial Actin Cytoskeleton by Directly Binding and Inducing a Low-Affinity State in MreB. Biochemistry. 2009;48(22):4852–7. doi: 10.1021/bi900014d. PubMed PMID: PMC3951351; PMCID: PMC2897334.

17. Gitai Z, Dye N, Shapiro L. An Actin-Like Gene can Determine Cell Polarity in Bacteria. Proceedings of the National Academy of Sciences of the United States of America. 2004;101(23):8643–8. doi: 10.1073/pnas.0402638101; PMCID: PMC423248.

18. Grinnell A, Sloan R, Morgenstein RM. Cell Density-dependent Antibiotic Tolerance to Inhibition of the Elongation Machinery Requires Fully Functional PBP1B. Commun Biol. 2022;5(1):107. Epub 2022/02/05. doi: 10.1038/s42003-022-03056-x. PubMed PMID: 35115684; PMCID: PMC8813938.

19. Barton B, Grinnell A, Morgenstein RM. Disruption of the MreB Elongasome Is Overcome by Mutations in the Tricarboxylic Acid Cycle. Front Microbiol. 2021;12:664281. Epub 2021/05/11. doi: 10.3389/fmicb.2021.664281. PubMed PMID: 33968001; PMCID: PMC8102728.

20. Baba T, Ara T, Hasegawa M, Takai Y, Okumura Y, Baba M, Datsenko KA, Tomita M, Wanner BL, Mori H. Construction of Escherichia coli K-12 in-frame, Single-gene Knockout Mutants: the Keio Collection. Mol Syst Biol. 2006;2:2006.0008-2006.0008. doi: 10.1038/msb4100050. PubMed PMID: PMC1681482.

21. Uehara T, Parzych KR, Dinh T, Bernhardt TG. Daughter Cell Separation is Controlled by Cytokinetic Ring-activated Cell Wall Hydrolysis. EMBO J. 2010;29(8):1412–22. Epub 2010/03/20. doi: 10.1038/emboj.2010.36. PubMed PMID: 20300061; PMCID: PMC2868575.

22. Uehara T, Dinh T, Bernhardt TG. LytM-domain Factors are Required for Daughter Cell Separation and Rapid Ampicillin-induced Lysis in Escherichia coli. J Bacteriol. 2009;191(16):5094–107. Epub 2009/06/16. doi: 10.1128/jb.00505-09. PubMed PMID: 19525345; PMCID: PMC2725582.

23. Cho H, Uehara T, Bernhardt TG. Beta-lactam Antibiotics Induce a Lethal Malfunctioning of the Bacterial Cell Wall Synthesis Machinery. Cell. 2014;159(6):1300–11. PubMed PMID: 25480295; PMCID: PMC4258230.

24. Govindarajan S, Amster-Choder O. The Bacterial Sec System is Required for the Organization and Function of the MreB Cytoskeleton. PLoS Genet. 2017;13(9):e1007017. Epub 2017/09/26. doi: 10.1371/journal.pgen.1007017. PubMed PMID: 28945742; PMCID: PMC5629013.

25. Bernhardt TG, de Boer PA. Screening for Synthetic Lethal Mutants in Escherichia coli and Identification of EnvC (YibP) as a Periplasmic Septal Ring Factor with Murein hydrolase activity. Mol Microbiol. 2004;52(5):1255–69. Epub 2004/05/29. doi: 10.1111/j.1365-2958.2004.04063.x. PubMed PMID: 15165230; PMCID: PMC4428336.

26. Mueller EA, Iken AG, Ali Öztürk M, Winkle M, Schmitz M, Vollmer W, Di Ventura B, Levin PA. The Active Repertoire of Escherichia coli Peptidoglycan Amidases Varies with Physiochemical Environment. Mol Microbiol. 2021;116(1):311–28. Epub 2021/03/06. doi: 10.1111/mmi.14711. PubMed PMID: 33666292; PMCID: PMC8295211.

27. Heidrich C, Ursinus A, Berger J, Schwarz H, Höltje JV. Effects of Multiple Deletions of Murein Hydrolases on Viability, Septum Cleavage, and Sensitivity to Large Toxic Molecules in Escherichia coli. J Bacteriol. 2002;184(22):6093–9. Epub 2002/10/26. doi: 10.1128/jb.184.22.6093-6099.2002. PubMed PMID: 12399477; PMCID: PMC151956.

28. Yakhnina AA, McManus HR, Bernhardt TG. The Cell Wall Amidase AmiB is Essential for Pseudomonas aeruginosa Cell Division, Drug Resistance and Viability. Mol Microbiol. 2015;97(5):957–73. Epub 2015/06/03. doi: 10.1111/mmi.13077. PubMed PMID: 26032134; PMCID: PMC4646093.

29. Ize B, Stanley NR, Buchanan G, Palmer T. Role of the Escherichia coli Tat Pathway in Outer Membrane Integrity. Mol Microbiol. 2003;48(5):1183–93. Epub 2003/06/06. doi: 10.1046/j.1365-2958.2003.03504.x. PubMed PMID: 12787348.

30. Hara H, Narita S, Karibian D, Park JT, Yamamoto Y, Nishimura Y. Identification and Characterization of the Escherichia coli envC Gene Encoding a Periplasmic Coiled-coil Protein with Putative Peptidase Activity. FEMS Microbiol Lett. 2002;212(2):229–36. doi: 10.1111/j.1574-6968.2002.tb11271.x.

31. Weatherspoon-Griffin N, Zhao G, Kong W, Kong Y, Morigen Andrews-Polymenis H, McClelland M, Shi Y. The CpxR/CpxA Two-component System Up-Regulates Two Tat-dependent Peptidoglycan Amidases to Confer Bacterial Resistance to Antimicrobial Peptide. The Journal of biological chemistry. 2011;286(7):5529–39. Epub 2010/12/13. doi: 10.1074/jbc.M110.200352. PubMed PMID: 21149452; PMCID: PMC3037666.

32. Kundig W, Ghosh S, Roseman S. PHOSPHATE BOUND TO HISTIDINE IN A PROTEIN AS AN INTERMEDIATE IN A NOVEL PHOSPHO-TRANSFERASE SYSTEM. Proc Natl Acad Sci U S A. 1964;52(4):1067–74. Epub 1964/10/01. doi: 10.1073/pnas.52.4.1067. PubMed PMID: 14224387; PMCID: PMC300396.

33. Irnov I, Wang Z, Jannetty ND, Bustamante JA, Rhee KY, Jacobs-Wagner C. Crosstalk Between the Tricarboxylic Acid Cycle and Peptidoglycan Synthesis in Caulobacter crescentus Through the Homeostatic Control of α-ketoglutarate. PLoS Genet. 2017;13(8):e1006978. Epub 2017/08/23. doi: 10.1371/journal.pgen.1006978. PubMed PMID: 28827812; PMCID: PMC5578688.

34. Vadia S, Tse JL, Lucena R, Yang Z, Kellogg DR, Wang JD, Levin PA. Fatty Acid Availability Sets Cell Envelope Capacity and Dictates Microbial Cell Size. Current biology : CB. 2017;27(12):1757-67.e5. Epub 2017/06/08. doi: 10.1016/j.cub.2017.05.076. PubMed PMID: 28602657; PMCID: PMC5551417.

35. Westfall CS, Levin PA. Comprehensive Analysis of Central Carbon Metabolism Illuminates Connections Between Nutrient Availability, Growth Rate, and Cell Morphology in Escherichia coli. PLoS Genet. 2018;14(2):e1007205. Epub 2018/02/13. doi: 10.1371/journal.pgen.1007205. PubMed PMID: 29432413; PMCID: PMC5825171.

36. van den Ent F, Amos LA, Lowe J. Prokaryotic Origin of the Actin Cytoskeleton. Nature. 2001;413(6851):39–44.

37. Deutscher J, Francke C, Postma PW. How Phosphotransferase System-related Protein Phosphorylation Regulates Carbohydrate Metabolism in Bacteria. Microbiol Mol Biol Rev. 2006;70(4):939–1031. Epub 2006/12/13. doi: 10.1128/mmbr.00024-06. PubMed PMID: 17158705; PMCID: PMC1698508.

38. Lopian L, Elisha Y, Nussbaum-Shochat A, Amster-Choder O. Spatial and Temporal Organization of the E. coli PTS Components. EMBO J. 2010;29(21):3630–45. Epub 2010/10/07. doi: 10.1038/emboj.2010.240. PubMed PMID: 20924357; PMCID: PMC2982763.

39. Dimitrova MN, Szczepanowski RH, Ruvinov SB, Peterkofsky A, Ginsburg A. Interdomain Interaction and Substrate Coupling Effects on Dimerization and Conformational Stability of Enzyme I of the Escherichia coli phosphoenolpyruvate:sugar phosphotransferase System. Biochemistry. 2002;41(3):906–13. doi: 10.1021/bi011801x.

40. Figge RM, Divakaruni AV, Gober JW. MreB, the Cell Shape-Determining Bacterial Actin Homologue, Co-Ordinates Cell Wall Morphogenesis in Caulobacter crescentus. Mol Microbiol. 2004;51(5):1321–32. doi: 10.1111/j.1365-2958.2003.03936.x. PubMed PMID: 14982627.

41. Yakhnina AA, Gitai Z. The Small Protein MbiA Interacts with MreB and Modulates Cell Shape in Caulobacter crescentus. Mol Microbiol. 2012;85(6):1090–104. doi: 10.1111/j.1365-2958.2012.08159.x.

42. Fenton AK, Gerdes K. Direct interaction of FtsZ and MreB is Required for Septum Synthesis and Cell Division in Escherichia coli. EMBO J. 2013;32(13):1953–65. Epub 2013/06/13. doi: 10.1038/emboj.2013.129. PubMed PMID: 23756461; PMCID: PMC3708099.

43. van der Ploeg R, Verheul J, Vischer NOE, Alexeeva S, Hoogendoorn E, Postma M, Banzhaf M, Vollmer W, den Blaauwen T. Colocalization and Interaction Between Elongasome and Divisome During a Preparative Cell Division Phase in Escherichia coli. Mol Microbiol. 2013;87(5):1074–87. doi: 10.1111/mmi.12150; PMCID: 23387922.

44. Yoshii Y, Niki H, Shiomi D. Division-site localization of RodZ is Required for Efficient Z ring Formation in Escherichia coli. Mol Microbiol. 2019;111(5):1229–44. Epub 2019/02/12. doi: 10.1111/mmi.14217. PubMed PMID: 30742332.

45. Campos M, Surovtsev IV, Kato S, Paintdakhi A, Beltran B, Ebmeier SE, Jacobs-Wagner C. A Constant Size Extension Drives Bacterial Cell Size Homeostasis. Cell. 2014;159(6):1433–46. doi: 10.1016/j.cell.2014.11.022. PubMed PMID: 25480302; PMCID: PMC4258233.

46. Taheri-Araghi S, Bradde S, Sauls JT, Hill NS, Levin PA, Paulsson J, Vergassola M, Jun S. Cell-size Control and Homeostasis in Bacteria. Current biology : CB. 2015;25(3):385–91. Epub 2014/12/24. doi: 10.1016/j.cub.2014.12.009. PubMed PMID: 25544609; PMCID: PMC4323405.

47. Peters NT, Dinh T, Bernhardt TG. A Fail-safe Mechanism in the Septal Ring Assembly Pathway Generated by the Sequential Recruitment of Cell Separation Amidases and Their Activators. J Bacteriol. 2011;193(18):4973–83. Epub 2011/07/19. doi: 10.1128/jb.00316-11. PubMed PMID: 21764913; PMCID: PMC3165665.

48. Ursinus A, van den Ent F, Brechtel S, de Pedro M, Höltje JV, Löwe J, Vollmer W. Murein (peptidoglycan) Binding Property of the Essential Cell Division Protein FtsN from Escherichia coli. J Bacteriol. 2004;186(20):6728–37. Epub 2004/10/07. doi: 10.1128/jb.186.20.6728-6737.2004. PubMed PMID: 15466024; PMCID: PMC522186.

49. Gerding MA, Liu B, Bendezú FO, Hale CA, Bernhardt TG, de Boer PA. Self-enhanced accumulation of FtsN at Division Sites and Roles for Other Proteins with a SPOR domain (DamX, DedD, and RlpA) in Escherichia coli Cell Constriction. J Bacteriol. 2009;191(24):7383–401. Epub 2009/08/18. doi: 10.1128/jb.00811-09. PubMed PMID: 19684127; PMCID: PMC2786604.

50. Boes A, Olatunji S, Breukink E, Terrak M. Regulation of the Peptidoglycan Polymerase Activity of PBP1b by Antagonist Actions of the Core Divisome Proteins FtsBLQ and FtsN. mBio. 2019;10(1). Epub 2019/01/10. doi: 10.1128/mBio.01912-18. PubMed PMID: 30622193; PMCID: PMC6325244.

51. García del Portillo F, de Pedro MA, Joseleau-Petit D, D’Ari R. Lytic Response of Escherichia coli Cells to Inhibitors of Penicillin-binding proteins 1a and 1b as a Timed Event Related to Cell Division. J Bacteriol. 1989;171(8):4217–21. Epub 1989/08/01. doi: 10.1128/jb.171.8.4217-4221.1989. PubMed PMID: 2666392; PMCID: PMC210193.

52. Colavin A, Hsin J, Huang KC. Effects of Polymerization and Nucleotide Identity on the Conformational Dynamics of the Bacterial Actin Homolog MreB. Proc Natl Acad Sci USA. 2014;111(9):3585–90. doi: 10.1073/pnas.1317061111; PMCID: PMC3948266

53. dos Remedios CG, Chhabra D, Kekic M, Dedova IV, Tsubakihara M, Berry DA, Nosworthy NJ. Actin Binding Proteins: Regulation of Cytoskeletal Microfilaments. Physiol Rev. 2003;83(2):433–73. Epub 2003/03/29. doi: 10.1152/physrev.00026.2002. PubMed PMID: 12663865.

54. Dye NA, Pincus Z, Fisher IC, Shapiro L, Theriot JA. Mutations in the Nucleotide Binding Pocket of MreB can Alter Cell Curvature and Polar Morphology in Caulobacter. Mol Microbiol. 2011;81(2):368–94. doi: 10.1111/j.1365-2958.2011.07698.x. PubMed PMID: PMC3137890; PMCID: PMC3137890.

55. Schaechter M, Maaloe O, Kjeldgaard NO. Dependency on Medium and Temperature of Cell Size and Chemical Composition During Balanced Grown of Salmonella typhimurium. J Gen Microbiol. 1958;19(3):592–606. Epub 1958/12/01. doi: 10.1099/00221287-19-3-592. PubMed PMID: 13611202.

56. Hill NS, Buske PJ, Shi Y, Levin PA. A Moonlighting Enzyme Links Escherichia coli Cell Size with Central Metabolism. PLoS Genet. 2013;9(7):e1003663. Epub 2013/08/13. doi: 10.1371/journal.pgen.1003663. PubMed PMID: 23935518; PMCID: PMC3723540.

57. Weart RB, Lee AH, Chien AC, Haeusser DP, Hill NS, Levin PA. A Metabolic Sensor Governing Cell Size in Bacteria. Cell. 2007;130(2):335–47. Epub 2007/07/31. doi: 10.1016/j.cell.2007.05.043. PubMed PMID: 17662947; PMCID: PMC1971218.

58. Waygood EB, Steeves T. Enzyme I of the phosphoenolpyruvate: sugar phosphotransferase system of Escherichia coli. Purification to Homogeneity and Some Properties. Can J Biochem. 1980;58(1):40–8. Epub 1980/01/01. doi: 10.1139/o80-006. PubMed PMID: 6992959.

59. Long CP, Au J, Sandoval NR, Gebreselassie NA, Antoniewicz MR. Enzyme I Facilitates Reverse Flux from Pyruvate to Phosphoenolpyruvate in Escherichia coli. Nat Commun. 2017;8:14316. Epub 2017/01/28. doi: 10.1038/ncomms14316. PubMed PMID: 28128209; PMCID: PMC5290146.

60. Monahan LG, Hajduk IV, Blaber SP, Charles IG, Harry EJ. Coordinating Bacterial Cell Division with Nutrient Availability: a Role for Glycolysis. mBio. 2014;5(3):e00935–14. Epub 2014/05/16. doi: 10.1128/mBio.00935-14. PubMed PMID: 24825009; PMCID: PMC4030479.

61. Deatherage DE, Barrick JE. Identification of Mutations in Laboratory-evolved Microbes from Next-generation Sequencing Data Using breseq. Methods Mol Biol. 2014;1151:165–88. Epub 2014/05/20. doi: 10.1007/978-1-4939-0554-6_12. PubMed PMID: 24838886; PMCID: PMC4239701.

62. Ursell T, Lee TK, Shiomi D, Shi H, Tropini C, Monds RD, Colavin A, Billings G, Bhaya-Grossman I, Broxton M, Huang BE, Niki H, Huang KC. Rapid, precise Quantification of Bacterial Cellular Dimensions Across a Genomic-scale Knockout Library. BMC Biology. 2017;15(1):17. doi: 10.1186/s12915-017-0348-8; PMCID: PMC5320674.

